# Oligogenic effects of 16p11.2 copy number variation on craniofacial development

**DOI:** 10.1101/540732

**Authors:** Yuqi Qiu, Thomas Arbogast, Sandra Martin Lorenzo, Honying Li, Shih C. Tang, Ellen Richardson, Oanh Hong, Shawn Cho, Omar Shanta, Timothy Pang, Christina Corsello, Curtis K. Deutsch, Claire Chevalier, Erica E. Davis, Lilia M. Iakoucheva, Yann Herault, Nicholas Katsanis, Karen Messer, Jonathan Sebat

## Abstract

A copy number variant (CNV) of 16p11.2, which encompasses 30 genes, is associated with developmental and psychiatric disorders, head size and body mass. The genetic mechanisms that underlie these associations are not understood. To elucidate the effects of genes on development, we exploited the quantitative effects of CNV on craniofacial structure in humans and model organisms. We show that reciprocal deletion and duplication of 16p11.2 have characteristic “mirror” effects on craniofacial features that are conserved in human, rat and mouse. By testing gene dosage effects on the shape of the mandible in zebrafish, we show that the distribution of effects for all individual genes is consistent with that of the CNV, and some combinations have non-additive effects. Our results suggest that, at minimum, one third of genes within the 16p11.2 region influence craniofacial development, and the facial gestalt of each CNV represents a product of 30 dosage effects.

**Highlights:**

- Reciprocal CNVs of 16p11.2 have mirror effects on craniofacial structure. Copy number is associated with a positive effect on nasal and mandibular regions and a negative effect on frontal regions of the face.
- Effects of CNV on craniofacial development in human are well conserved in rat and mouse models of 16p11.2 deletion and duplication.
- 7/30 genes each independently have significant effects on the shape of the mandible in zebrafish; these include *SPN, C16orf54, SEZ6L2, ASPHD1, TAOK2, INO80E* and *FAM57B*. Others (*MAPK3, MVP, KCTD13*) have detectable effects only in combination.
- Overexpression of 30 genes individually showed a distribution of effects that was skewed in the same direction as that of the full duplication, suggesting that specific facial features represent the net of all individual effects combined.

## Introduction

Recent technological advances in genomics have facilitated the discovery of scores of new genetic disorders that have a complex and variable clinical presentation (Malhotra and Sebat, 2012). Unlike Down syndrome (Roizen and Patterson) and Williams’s syndrome (Ewart et al.), which have a distinguishable constellation of clinical features and facial gestalts, these new genetic disorders are notable for not having a clear pattern of congenital anomalies or dysmorphic features (Nevado et al.). A major exemplar are the reciprocal CNVs of 16p11.2 (BP4-BP5, OMIM: 611913 and 614671). Deletion (Miller et al.) and duplication (D’Angelo et al.) of 30 genes are associated with variable degrees of cognitive impairment, epilepsy and psychiatric traits including autism spectrum disorder, psychiatric disorders. We and others have shown that the dosage of 16p11.2 has quantitative effects on development, in particular morphometric traits, such as head circumference (McCarthy et al.; Shinawi et al.) and body mass index (BMI) (D’Angelo et al.). Deletions are associated with greater head size and BMI while duplications are associated with smaller head size and BMI. In addition, a variety of craniofacial anomalies have been reported in a subset of cases (Bijlsma et al.; Rosenfeld et al.; Shinawi et al.), but a characteristic pattern of dysmorphic features has not been described. Thus, the influence of CNV on psychiatric and morphometric traits alike is complex, and the underlying genetic mechanisms are not understood.

Elucidating the genetic mechanisms through which CNVs influence development requires rigorous analysis of quantitative phenotype data in humans and the establishment of model systems in which the genetic mechanisms are conserved. Craniofacial development, in particular, is controlled by genetic mechanisms that are conserved across species (Schilling). The effect of genes on the human face is of interest, therefore, because craniofacial structure represents developmental phenotypes that are experimentally tractable in model organisms, and which could provide insights into disease mechanisms. Effects of 16p11.2 CNV on development of the brain and head have been reported in both mouse (Arbogast et al.; Horev et al.) and zebrafish (Golzio et al.), and multiple genes have been demonstrated to influence brain development including KCTD13, MAPK3 and MVP (Arbogast et al.; Escamilla et al.; Golzio et al.). We hypothesize that a precise morphometric characterization of patients could help to further illuminate how 16p11.2 genes influence embryonic development.

The application of 3D imaging provides detailed quantitative analysis of surface features, enabling more precise measurements of the shape of the head and face. Application of this approach has facilitated the finer characterization of genetic syndromes with characteristic craniofacial features (Hammond; Hammond et al.). Application of this technology in non-syndromic and complex genetic disorders has the potential to elucidate the effect of genes on craniofacial development. By three-dimensional image analysis of surface features in human, rat and mouse and the dissection of single gene effects in zebrafish, we show that the copy number of 16p11.2 has a strong effects on craniofacial structure that are conserved across species, and the facial features associated with each disorder are attributable to the oligogenic effects of multiple genes.

## Results

### Reciprocal deletion and duplication of 16p11.2 have mirror effects on craniofacial structure

3D morphometric facial imaging was performed on subjects with 16p11.2 duplications or deletions and controls recruited to the Simons VIP study (Simons VIP Consortium, see supplementary methods). The final dataset (N = 228, **Table S1**) included 45 with deletions, 44 with duplications and 139 familial non-carrier controls. A total of 24 landmarks were placed on each image and a “features”, defined as pairwise distances between landmarks, were normalized to the mean. Differences in deletion and duplication groups relative to controls were detected by linear regression controlling for fixed effects of age, head circumference, body mass index (BMI), sex, and ancestry principal components obtained from genetic data, with a random intercept allowed to account for within-family correlation.

Eighteen features differed significantly between groups at a family-wise error rate of 5% (**Fig 1B**, **Table S2**), and forty-five were significant at a Benjamini-Hochberg FDR correction of 5%. For 13 of the 18 significant features, deletion and duplication had effects that were opposite in direction (p= 0.048, one-sided binomial test). Consistent with the deletion and duplication having reciprocal effects, the deletion vs duplication effect sizes were negatively correlated for the 18 significant measures (p = <0.001, Pearson’s correlation = −0.77, **Fig 1A**).

**Figure 1.**
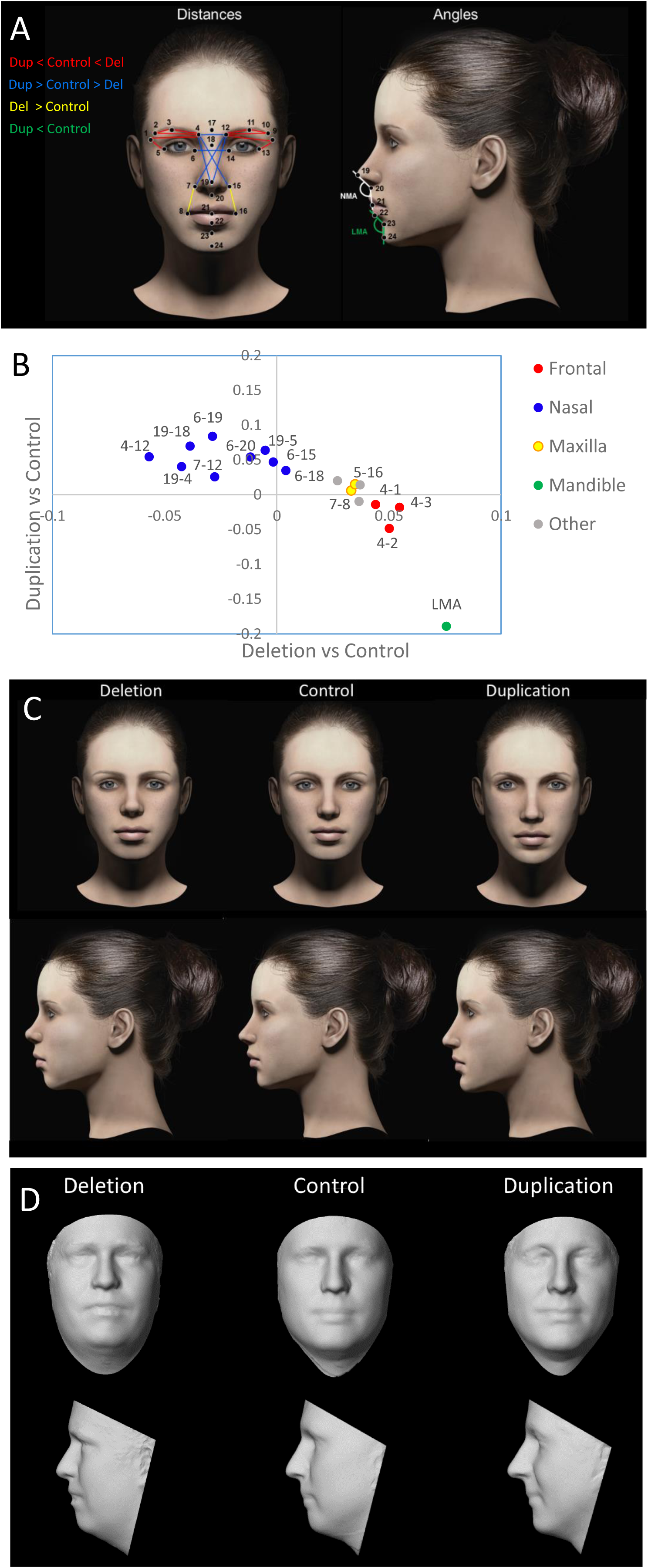
Differential effects of 16p11.2 copy number on dimensions of the frontal, nasal, maxillary and mandibular regions. **(A)** On each 3D facial image, 24 landmarks were placed and two angular measurements were calculated. After averaging symmetric distances, 156 distance measures were compared between the CNV and control groups. **(B)** 18 measures were significant after correction for a FWER < 5%. Regression coefficients for duplication vs control (y-axis) and deletion vs control (x-axis) show that reciprocal CNVs have reciprocal effects on growth of the major craniofacial processes. The category “Other” represents features that span multiple processes. The 14 most informative facial features based on LASSO selection are drawn in panel A and colored by facial region according to the legend. For clarity some nasal distances are excluded. **(C)** Facial features associated with deletion and duplication were visualized by adjusting the computer-generated model face according the observed effect sizes (from Panel B and **Table S2**). **(D)** The average surface topography was generated from multiple (>5) age-matched subjects with each genotype. Note that subtle differences in BMI are also apparent, however these effects are controlled for in the statistical analysis, and do not influence the feature selection.

Genetic effects were clustered in regions corresponding to major processes of craniofacial development (Frontonasal, Medial Nasal, Maxilla and Mandible, **Fig 1A**). Deletion of 16p11.2 was associated with significantly larger frontal (4-1, 4-2, 4-3, 12-9, 12-10, 12-11) and maxillary (7-8, 15-16, 5-16 and 13-8) dimensions and a shorter (18-19) and narrower nose (4-12, 4-15, 6-15, 7-12 and 7-14). By contrast the duplication was associated with opposite effects, including smaller frontal dimensions (4-2) and significantly wider nose and longer nasal bridge (18-19). Duplications were associated with a narrower LMA consistent with a more protrusive chin. A wider LMA was observed in deletion carriers, but the effect did not reach statistical significance in this comparison. Least absolute shrinkage and selection operator (LASSO) logistic regression was performed to select a parsimonious subset of 14 features which could best discriminate each genotype (**Fig. 1B**, **Table S2**).

Facial gestalts associated with the 16p11.2 deletion and duplication were visualized using computer-generated faces in which the features of a model face were adjusted according to the 14 differences described above, including the frontonasal and maxillary distances and the LMA (**Fig. 1C**, **Table S2)**. Dimensions were adjusted based on the percentage difference between CNV and control groups (defined as the effect size divided by the mean). Differences ranged from 1% to 12%.

To further visualize the facial gestalts of controls, deletion carriers and duplication carriers respectively, a 3D-model of each was generated by averaging of the surface topography of faces from multiple subjects (3dMDvultus version 2.5.0.1). The sample was limited to subjects ages 14-49 to avoid variability in facial features across development at young ages, and the sample was restricted to males for which there were sufficient numbers of age-matched subjects (>5) for all three genotypes (**Table S3**). A facial gestalt similar to that of the simulated faces was distinguishable in the average faces of adult male deletion and duplication carriers and control subjects, with the effects on the nose and chin being the most recognizable feature (**Fig. 1D**). Similar features were observed in the average faces of younger (age 8-11) and older (ages 18-50) subjects of both sexes (**Fig. S1**).

### Craniofacial characteristics distinguish 16p11.2 deletion and duplication carriers from controls

Based on Linear Discriminate Analysis (LDA) of craniofacial features, genotypes could be separated into clusters, with better separation for younger subjects (**Fig. 2**). The LDA model achieved a total correct classification rate of 0.78 on the full sample; reflecting the considerable overlap between the genotypes (**Fig. 2A**). Genotype was classified more accurately by LDA when restricted to younger (age 3-20) subjects, with total correct classification 0.84. The predictive accuracy of the LDA model was confirmed by leave-one-out cross validation of the full sample which gave specificities of 0.88 and 0.93 and sensitivities of 0.48 and 0.42 for deletion and duplication respectively. When restricted to younger subjects, specificities were 0.88 and 0.87 and sensitivities were 0.72 and 0.52 for deletion and duplication respectively.

**Figure 2.**
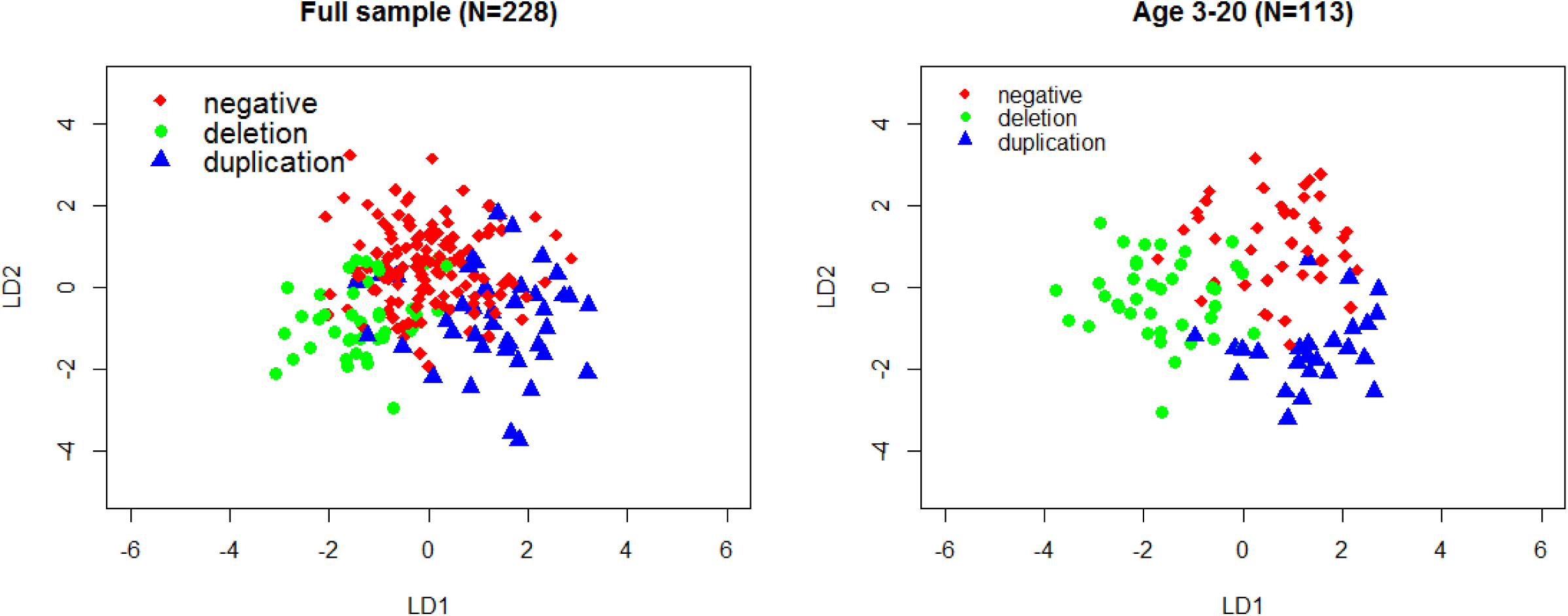
Classification of 16p11.2 genotype based on facial features. Discriminate coefficients based on features that were significant at FDR < 0.05 can distinguish the subjects based on genotype, with better discrimination for younger subjects (age ≤ 20 years). The linear model was controlled for age, head circumference, body mass index (BMI), sex, and ancestry principal components. LDA was applied to subjects for which the above demographic information was complete (N= 220 for the full sample and 107 for the younger group).

These results demonstrate that deletion and duplication carriers have combinations of facial features that are distinctive for each group. However, the substantial overlap between the faces of CNV carriers and controls is consistent with many subjects having a non-syndromic appearance that is not characterized by gross anomalies. Examination of group differences on each of the individual distances confirms that deletion and duplication groups do not represent outliers on any single measure. (**Fig. S2**).

### Differential effects CNV on craniofacial structure are recapitulated in rat and mouse models of 16p11.2

Rodent models of 16p11.2 deletion and duplication exhibit a variety of behavioral traits (Arbogast et al.; Horev et al.; Yang et al.). However, the direct relevance of these phenotypes to the human condition is uncertain. Similarly, the analysis of anthropometric traits in model organisms has been confounded by growth retardation that is observed in some mouse models (Arbogast et al.; Horev et al.; Yang et al.). We theorized that the cranial skeleton might represent an aspect of vertebrate development that is sufficiently conserved to serve as surrogate traits for genetic dissection of 16p11.2 CNV. To that end, we pursued quantitative analyses of the cranial skeleton from rat and mouse models of the 16p11.2 deletion and duplication (Arbogast et al., 2016).

Rat deletion and duplication models were generated by CRISPR/Cas9 genome editing of the syntenic region, and computed tomography (CT) scans were obtained from a cohort of 75 rats. In addition, CT scans of mouse lines from Arbogast et al. were obtained from a cohort of 26 mice (see supplementary methods). For each subject, a set of 19 landmarks were placed delineating the major craniofacial processes, and features were compared between the CNV models and matched controlled using linear regression. Results of all univariate tests are described in **Table S4**.

CNV had a significant effect on craniofacial structure in rat with strong mirror effects across all features between the deletion and duplication models (r = −0.56, P < 0.001, **Fig. S3**). A total of 52 features were significantly associated with genotype (FDR < 0.05, **Fig 3A**). By labeling features according to their respective craniofacial regions, we observe that the deletion was associated with larger frontal regions (e.g. 9-2, 9-3, 9-6 and 9-7, **Fig. 3B**) and smaller nasal (3-7, 3-8, 7-7 and 4-8) and mandibular (MW) regions, while the opposite effects were associated with the duplication. These results are consistent with the patterns that were observed in human.

**Figure 3.**
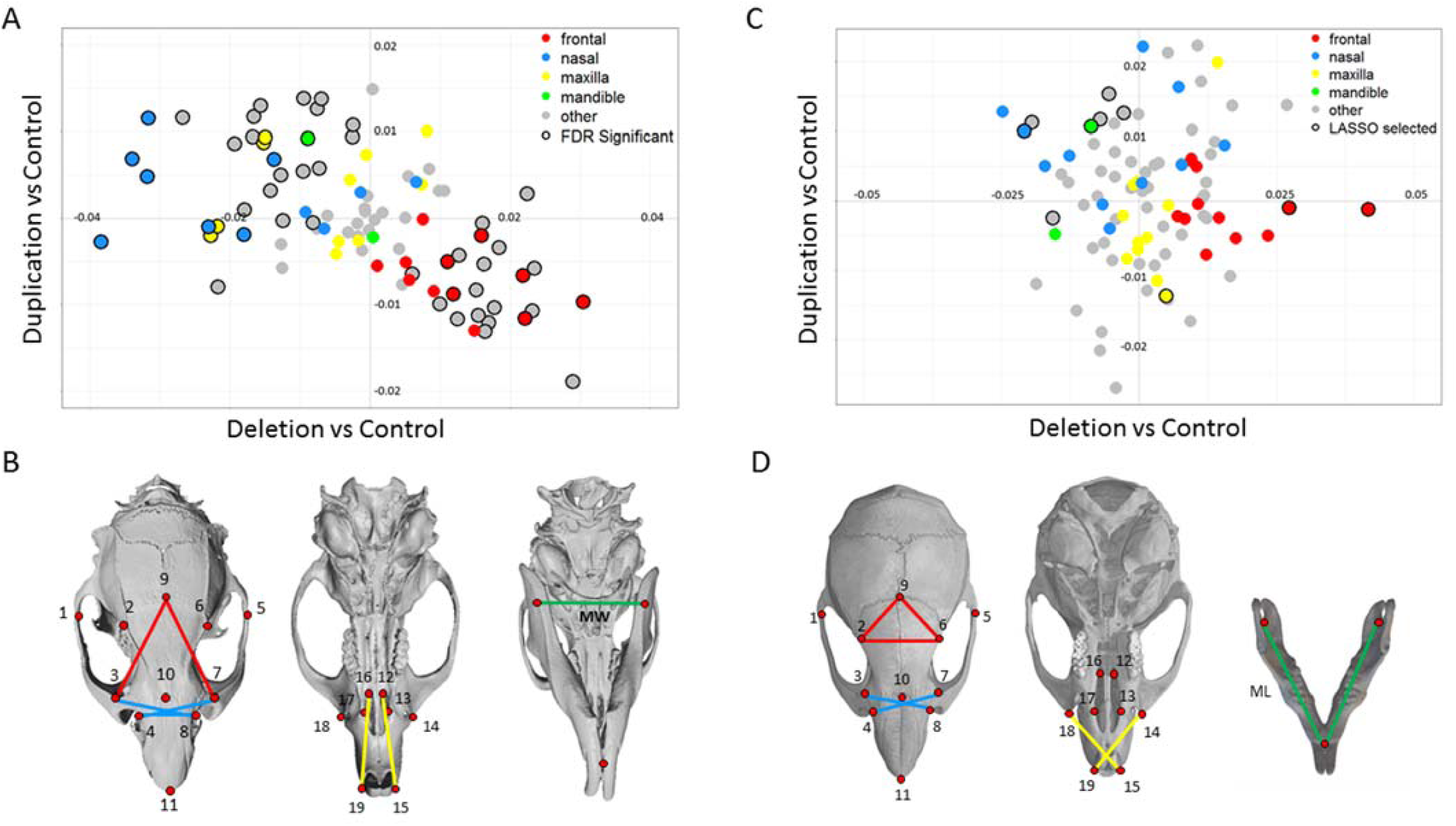
Validation of mirror craniofacial effects in rat and mouse models of 16p11.2 deletion and duplication. All pairwise distances were analyzed for nineteen landmarks on the dorsal skull and three on the mandible as shown here and in Table S4. Distances are colored according to craniofacial region using the same scheme as in figure 1. Distances that span multiple craniofacial processes are denoted as “other”. ML = Mandibular length, MW = Mandibular Width. (**A**) In the rat models, 52 individual features differed significantly by genotype. Regression coefficients for the duplication deletion show significant mirror effects. (**B**) Informative features were identified by LASSO selection, and features that correspond to a specific facial process in rat are shown. (**C**) In the mouse models, twelve craniofacial measures that discriminated mutant and control groups were selected by LASSO. Regression coefficients of these features show mirror effects of deletion and duplication similar to those in human and rat. (**D**) Features that correspond to specific facial processes in mouse.

Overall the effects of the deletion in mouse were similar to those in rat with effect sizes across the face being significantly correlated between species (r = 0.50, p <0.0001, **Fig S10**). The effects of the duplication in mouse did not correlate with those in rat and did not exhibit a strong mirror effect relative to the deletion across all features (p = 0.59), consistent with the duplication having a comparatively modest effect in this mouse line. Mouse craniofacial features that differed between deletion and duplication lines, however, did show mirror patterns similar to those in rat and human (**Fig. 3C**). For the most informative features that were selected by LASSO regression, deletion mice had larger frontal (9-2, 9-6 and 2-6) and maxillary (19-14 and 15-18) distances and smaller nasal (7-1, 8-1, 8-3 and 10-1) and mandibular (ML) distances, which were similar to the effects observed in human (sign test P = 0.004), whereas the duplication mouse model had reciprocal effects on the same features (sign test P = 0.004).

### Craniofacial features associated with 16p11.2 CNVs are attributable to multiple genes

To assess with more granularity the influence of the 16p11.2 BP4-BP5 genes on facial structure, we tested their effects on specific craniofacial features that could be measured by *in vivo* imaging of zebrafish larvae. Protrusion of the lower jaw was measured using the ceratohyal arch angle (CHA), where a smaller angle corresponds to a more protrusive jaw and a wider angle corresponds to a receding jaw (**Fig. 4A)**. Dimensions of the frontonasal region were measured using the Frontonasal area (FNA) and interocular distance (IOD) (**Fig. S4A**), however we are unable to capture separate frontal and nasal measurements in zebrafish analogous to those in rodent and in human.

**Figure 4.**
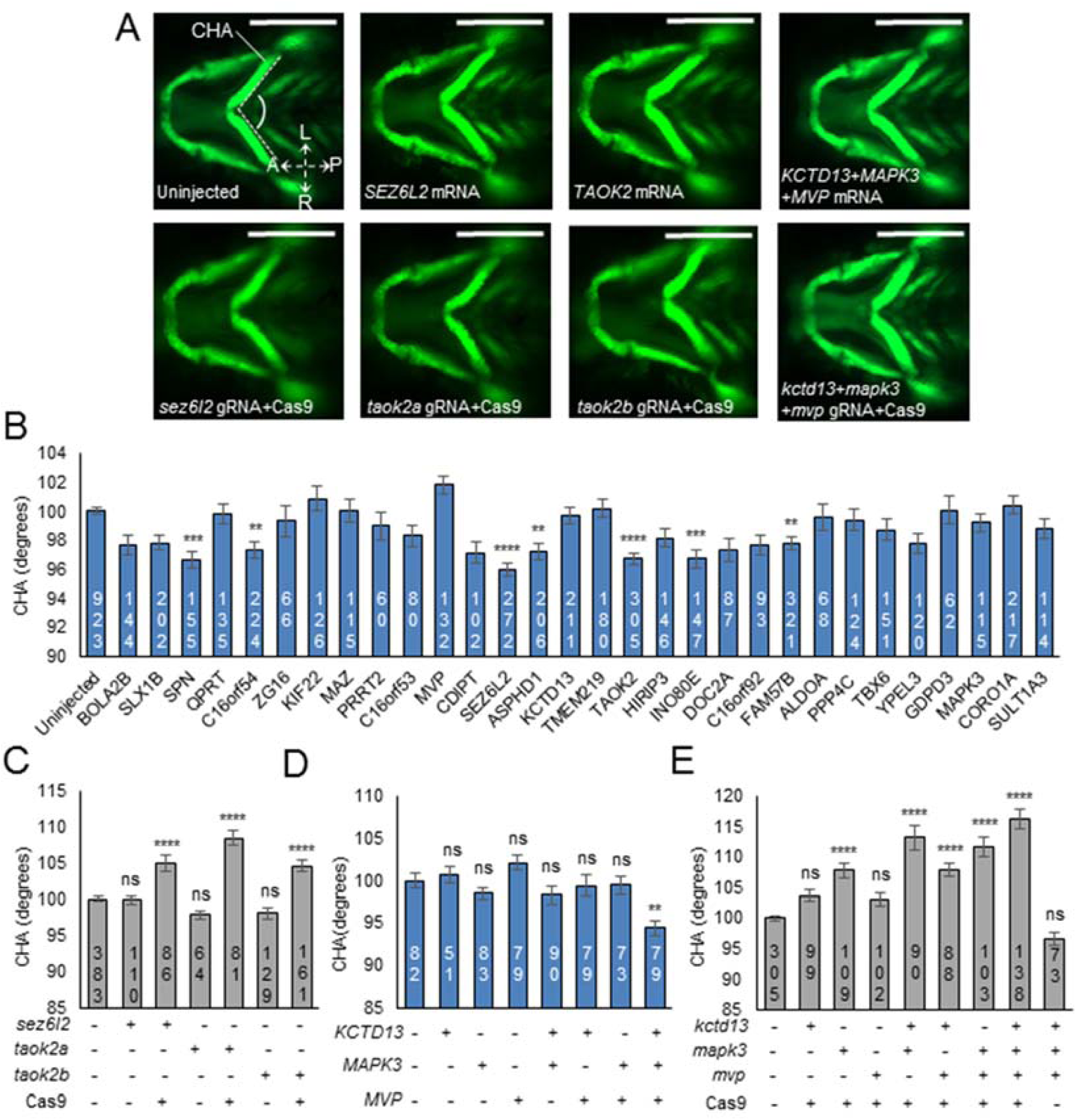
*In vivo* modeling of the 16p11.2 CNV implicates single gene drivers and epistatic effects influencing cartilage structures in the zebrafish pharyngeal skeleton. (**A**) Representative ventral images of *-1.4col1a1:egfp* zebrafish larvae at 3 days post-fertilization (dpf). Orientation arrows indicate anterior (A), posterior (P), left (L) and right (R). Scale bar, 300 µm. (**B**) Quantitative assessment of the CHA of larvae injected with single human mRNAs for the 30 genes located in the 16p11.2 BP4-BP5 region. Images were measured as shown in (**A**) (angle between dashed lines). Seven transcripts induced a significant reduction in CHA. Dosage: 12.5 pg for *KIF22* and *PPP4C*; 50 pg for all other genes. (**C**) Quantitative assessment of the CHA of F0 mutant batches injected with single combinations of each of *sez6l2, taok2a*, and *taok2b* gRNAs with or without Cas9. Dosage: 50 pg gRNA and 200 pg Cas9 protein. (**D**) Quantitative assessment of the CHA of larvae injected with single or equimolar combinations of human *KCTD13, MAPK3*, and *MVP* mRNAs. Dosage: 50 pg. (**E**) Quantitative assessment of the CHA of F0 mutant batches injected with single or equimolar combinations of *kctd13, mapk3*, and *mvp* gRNAs with or without Cas9. Dosage: 50 pg gRNA and 200 pg Cas9 protein. Number of larvae measured for each condition are indicated at the base of each bar in the graphs. The data are represented as the mean ± standard error of the mean (s.e.m.); ns=not significant; \*\**P*<0.01, \*\*\**P*<0.001 and \*\*\*\**P*<0.0001 vs uninjected controls. Tukey’s test was applied following a significant one-way ANOVA.

We first tested the overexpression of all 30 genes in the 16p11.2 region, focusing on the lower jaw phenotype which is more directly analogous to the phenotypes in human and rodent. We found that several genes had significant effects on CHA, including *SPN, C16orf54, SEZ6L2, ASPHD1, TAOK2, INO80E* and *FAM57B* (**Fig. 4B**), The genes inducing the most significant phenotypes included *SEZ6L2* (4° decrease in CHA versus controls; p<0.0001) and *TAOK2* (3° decrease in CHA versus controls; p<0.0001; **Fig. 4A,B**).. Effects for all seven transcripts were associated with a narrower CHA compared to controls, consistent with the protruding lower jaw that is associated with the duplication in human and rodent The distribution of effects for all 30 genes (95% CI = 98.1-99.1) was significantly lower than the distribution in controls (95% CI = 99.5-100.5), consistent with the effect of the duplication. We evaluated the effects of ablating endogenous zebrafish *sez6l2, taok2a*, and *taok2b* using CRISPR/Cas9 genome editing, and confirmed that the reciprocal loss of these genes results in a reciprocal increase of the CHA in comparison to controls (*sez6l2* gRNA+Cas9 versus controls, 5° increase in CHA, p<0.0001; *taok2a* gRNA+Cas9 versus controls, 8° increase in CHA, p<0.0001; *taok2b* gRNA+Cas9 versus controls, 5° increase in CHA, p<0.0001; **Fig. 4A,C**), consistent with the effect of the deletion.

We showed previously using zebrafish models that overexpression of *KCTD13* individually and in combination with *MAPK3* and *MVP* led to a decrease in head width (Golzio et al., 2012), and knockdown of *kctd13* exhibited mirror effects, a pattern consistent with the human phenotype of the 16p11.2 CNV. We tested overexpression and CRISPR/Cas9 F0 mutants of *KCTD13, MAPK3*, and *MVP* individually and in combinations of two or three genes. Overexpression of the three mRNAs individually did not have a significant effect on CHA, but injection of all three transcripts combined resulted in a significant 6° decrease in CHA relative to controls (Tukey’s p<0.01; **Fig. 4A, D**). Mutants with reciprocal loss of *mapk3* displayed an increased CHA (**Fig. 4E**) and the three gene combination resulted in a 16° CHA increase (Tukey’s p<0.0001). Thus mirror effects of these genes parallel those that are observed in human. We evaluated the body length of larvae injected with a combination of the three gRNAs and Cas9 and found no growth retardation compared to controls, supporting further the specificity of the cartilage phenotypes (**Fig. S5**). For Frontonasal area (FNA) and Interocular distance (IOD), significant effects were also observed with combinations of two or three genes (**Fig. S4**). Genome editing was associated with reduction in FNA (**Fig. S4A-C**), and gene overexpression was associated with increase in IOD (**Fig. S4D, E**), results that parallel the effect of the deletion and duplication on nasal regions in human. Evidence for a synergistic effect of MAPK3 in combination with MVP or KCTD13 was observed for dimensions of the frontonasal region but not the mandible (**Table S5**). Other combinations were consistent with additive effects (p=0.99 for additive ANOVA model compared to fully parameterized model). Together, our *in vivo* experiments performed in zebrafish suggest that facial features that are associated with CNV are under the influence of a substantial proportion of 16p11.2 genes, including some that have non-additive effects.

## Discussion

Here we show that reciprocal CNVs of the 16p11.2 BP4-BP5 region have mirror effects on craniofacial development. Deletion and duplication of 16p11.2 are each associated with facial features that are distinctive but both groups overlap with the variability observed in the general population. Dosage of 16p11.2 was associated with a positive effect on nasal and mandibular regions and a negative effect on the frontal regions.

The principal value of the 16p11.2 CNV facial phenotypes are, not as a clinical diagnostic markers, but as a model for studying the genetic mechanisms through which CNVs influence complex traits. Here we show that mirror effects of CNV on facial features are well conserved in rat and mouse models of 16p11.2, and the effects of gene dosage on a specific feature (shape of the mandible) can be further modeled in zebrafish. The craniofacial phenotype of 16p11.2 thus represents the first set of traits for which a genetic mechanisms is conserved across model systems.

By dissection of individual gene effects in zebrafish we show that mirror facial phenotypes of CNV are attributable to multiple genes within the region. Significant effects were observed for seven genes when overexpressed individually and for additional genes (MAPK3, MVP and KCTD13) when overexpressed in combination, thus at least one third of genes may influence the shape of the mandible The distribution of effects for all 30 gene overexpression tests was negative, consistent with the effect for the full duplication. Our results suggest that the net effect of the large CNV on specific developmental features consists of a combination of 30 individual gene effects. We find some evidence for synergistic effects when multiple genes are expressed in combination; however, we are not able to assess whether the overall effect is explained predominantly by additive effects or epistasis without a model of the full CNV in zebrafish.

The genes that have the greatest effects on shape of the mandible were *SPN, C16orf54, SEZ6L2, ASPHD1, TAOK2, INO80E and FAM57B*. These genes were not clearly distinguishable from the other 23 based on their levels of expression in the developing face (**Table S6**). However, some of these genes have been shown previously to be associated with alterations in head and brain size, such as *TAOK2* (Richter et al.) and *FAM57B* (McCammon et al.). These and other genes within the region function as regulators of cell proliferation and embryonic development (Khosravi-Far et al.). Notably, *TAOK2* is a regulator of MAP Kinase (MAPK) signaling (Chen et al.), which is a commonality among multiple 16p11.2 genes, including *MAPK3* which encodes the Extracellular Receptor Kinase 1 (ERK1) (Meloche and Pouyssegur) and *MVP* (Scheffer et al.), which complexes with ERK2 (Kolli et al.) and regulates ERK signaling (Kim et al.). This pathway-level convergence highlights MAPK signaling as a potential driver of craniofacial effects that are observed in this study.

The craniofacial features that are associated with the deletion of 16p11.2, including macrocephaly, broad forehead, and underdeveloped nose and chin (micrognathia), bare some similarity to features of monogenic disorders that are caused by mutations in components of RAS/MAPK signaling, such as Noonan (Bhambhani and Muenke) and Cardiofaciocutaneous (CFC) syndromes (Rauen). Similar craniofacial anomalies are also observed in mouse embryos with conditional disruption of MAPK signaling in neural crest cells (Parada et al.). Common facial features between 16p11.2 deletions and a subset of these other syndromes is intriguing and suggests that dysregulation of RAS/MAPK signaling might affect embryonic patterning in similar ways in 16p11.2 microdeletion syndrome and in the family disorders known as the “rasopathies” (Araki et al.).

An oligogenic mechanism is unlikely to be unique to the to the 16p11.2 locus. Rather an oligogenic model may apply in general to the effect of large CNVs on complex traits. For example, the polygenic contribution to height appears to be distributed across a large proportion of the genome (Boyle et al.; Liu et al.). The same is likely to be true for other anthropometric and cognitive traits such as facial features, body mass and IQ. In principle, haploinsufficiency of 30 adjacent genes may exert a distribution of effects across a variety of traits, and the features that are most prominent for a particular disorder could be those traits for which the sum of gene dosage effects across the CNV deviates significantly from the genome-wide genetic load.

Previous studies have found evidence that multiple genes within the 16p11.2 region impact various aspects of development in zebrafish (McCammon et al.) and drosophila (Iyer et al.). However, a major limitation has been a lack of validation of these phenotypes as models of the human disorder. The reciprocal craniofacial phenotypes that we observe are, to our knowledge, the only human 16p11.2-associated traits that are reproducible across multiple model organisms, both in magnitude and direction of effect. Knowledge of the influence of 16p11.2 deletion and duplication on craniofacial development could serve as a guide for how these genetic disorders influence embryonic patterning more broadly, including regional patterning of the brain (Chang et al.; Owen et al.; Owen et al.; Qureshi et al.). Further studies of the oligogenic effects described here could provide insights into mechanisms underlying cognitive impairments of these genetic disorders.

### Experimental Procedures

#### 3D Morphometric Analysis of Simons VIP subjects

Images of Simons VIP subjects were acquired using the 3dMDtrio system (http://www.3dmd.com/3dmd-systems/#trio) and were landmarked according to Farkas standards (Farkas) blind to genotype. Additional landmarks were placed to capture frontal dimensions including landmarks 2 (lateral brow) and 4 (medial brow). A total of 24 landmarks were placed. Quantitative pairwise distances between landmarks were calculated using the 3DMD software (3dMDvultus version 2.5.0.1). Symmetric distances were averaged and normalized to the overall size of the face, yielding 156 facial measurements and two angular measurements, the nasomental (NMA) and labiomental (LMA) angles respectively. We used a series of linear mixed-effects models to test for the effect of the deletion and the duplication separately on each facial measure. Linear regression was controlled for fixed effects of age, head circumference, body mass index (BMI), sex, and ancestry principal components obtained from genetic data, with a random intercept allowed to account for within-family correlation.

To investigate further the extent to which the 16p11.2 genotype can be distinguished based on craniofacial features, we performed Linear Discriminate Analysis (LDA) using the 45 distances significant at FDR <0.05 (**Fig. 2**), for the total sample and a subset restricted to younger subjects (age 3-20). Statistical analysis controlled for age, head circumference, body mass index (BMI), sex, and ancestry principal components. Leave-one-out cross validation was performed twice to calculate misspecification rates for both full sample and younger sample.

#### Computed tomography analysis of 16p11.2 deletion and duplication rodent models

Quantitative analysis of skull morphometry in rodent models of the 16p11.2 deletion and duplication was performed using a dataset of CT scans collected by Arbogast et al (Arbogast et al.) and completed here with a CT dataset for the rat models (see supplementary materials). The cohort of 75 rats consisted of 23 Del/+ (9 male and 14 female), 26 Dup/+ (13 male and 13 female) and 26 +/+ littermates (13 male and 13 female). The mouse cohort consisted of female deletion or duplication lines (aged 13 weeks) paired with the same number of wild-type female littermates, including 10 deletion and 8 duplication pairs obtained from more than 5 independent breeder pairs to minimize inbreedism.

Following CT scans of all rodents, nineteen landmarks were placed to capture the dimensions of the frontal, nasal and maxillary regions. Centroids of multiple landmarks were determined to mark three positions on the mandible (**Fig. S6**). For each skull, pairwise distances were normalized to the overall geometric mean distance, and the most significant differences between mutant and control groups were identified by linear regression at an unadjusted significance of 5%.

#### Testing the effects of genes on craniofacial features in a Zebrafish model

To model the 16p11.2 BP4-BP5 duplication, we overexpressed all 30 genes of the CNV individually and evaluated the shape of the mandible in a transgenic zebrafish model (-*1.4col1a1:egfp*) in which GFP marks the developing cartilage (Kague et al. 2012). In addition, we investigated pairwise and three-way gene interactions for *MAPK3, MVP*, and *KCTD13* that have been reported previously to impact head size (Golzio et al 2011). To model the reciprocal deletion, we performed CRISPR/Cas9 genome editing of *sez6l2, taok2a*, and *taok2b* individually; we generated F0 mutants for *mapk3, mvp*, and *kctd13* individually and in pairwise and three-way combinations. We used CHOPCHOP (Labun et al., 2016) to identify guide (g)RNAs targeting coding sequence and PCR primers used to determine efficiency of CRISPR/Cas9 genome editing (**Fig. S7-12**, **Table S7**). Measurements of the dorsal zebrafish head were performed by determining the interocular distance (IOD) and the frontonasal area (FNA).

## Acknowledgements

This study was funded by grants to J.S. from the Simons Foundation (SFARI #178088) and the NIMH (*new grant number pending); a grant to E.E.D. from the NIMH (MH106826); a NIMH Silvio O. Conte Center grant to N.K. (P50MH094268); a grant to Y.H. from the French Foundation for Rare Diseases; and grants to L.M.I. from the Simons Foundation (SFARI #345469) and NIMH (MH104766). We are grateful to all of the families at the participating Simons Variation in Individuals Project (Simons VIP) sites, as well as the Simons VIP Consortium. We appreciate obtaining access to 3D human imaging and genetic data from SFARI. Approved researchers can obtain the Simons VIP datasets described in this study (https://www.sfari.org/resource/simons-vip/) by applying at https://base.sfari.org. We are grateful to the CELPHEDIA/TEFOR French National infrastructure and especially to Severine Menoret, for helping us to obtain the 16p11.2 rat models that are described here.

## Supplementary Materials

### Supplementary Figures

**Figure S1.**
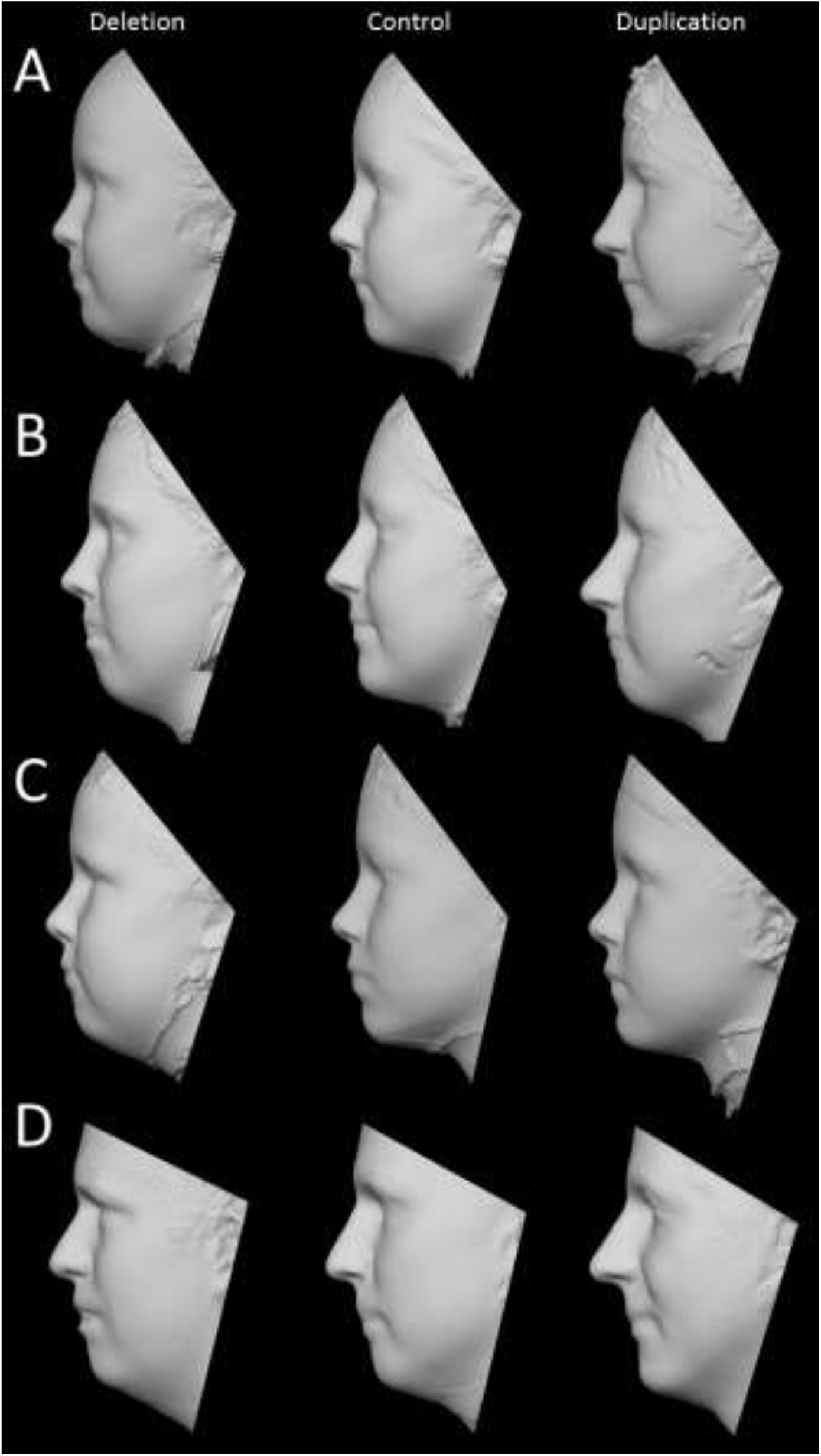
3D models of deletion, control and duplication groups generated by averaging of the surface topography of faces from multiple subjects. Separate models were constructed for adults and children of each sex. The number and mean age of subjects in each group is listed below. **(A) Female children subjects:** Deletion n= 7; mean age= 9.15 years; Control n= 8, mean age= 9.92 years; and Duplication n= 5, mean age= 12.73 year **(B) Female adult subjects:** Deletion n= 4; mean age= 20.13 years; Control n= 8, mean age= 23.71 years; and Duplication n= 5, mean age= 23.25 years **(C) Male children subjects:** Deletion n= 7; mean age= 8.90 years; Control n= 9, mean age= 9.27 years; and Duplication n= 9, mean age= 9.20 years **(D) Male adult subjects:** Deletion n= 5; mean age= 25.53 years; Control n= 10, mean age= 36.59 year; and Duplication n= 5, mean age= 36.48 years

**Figure S2.**
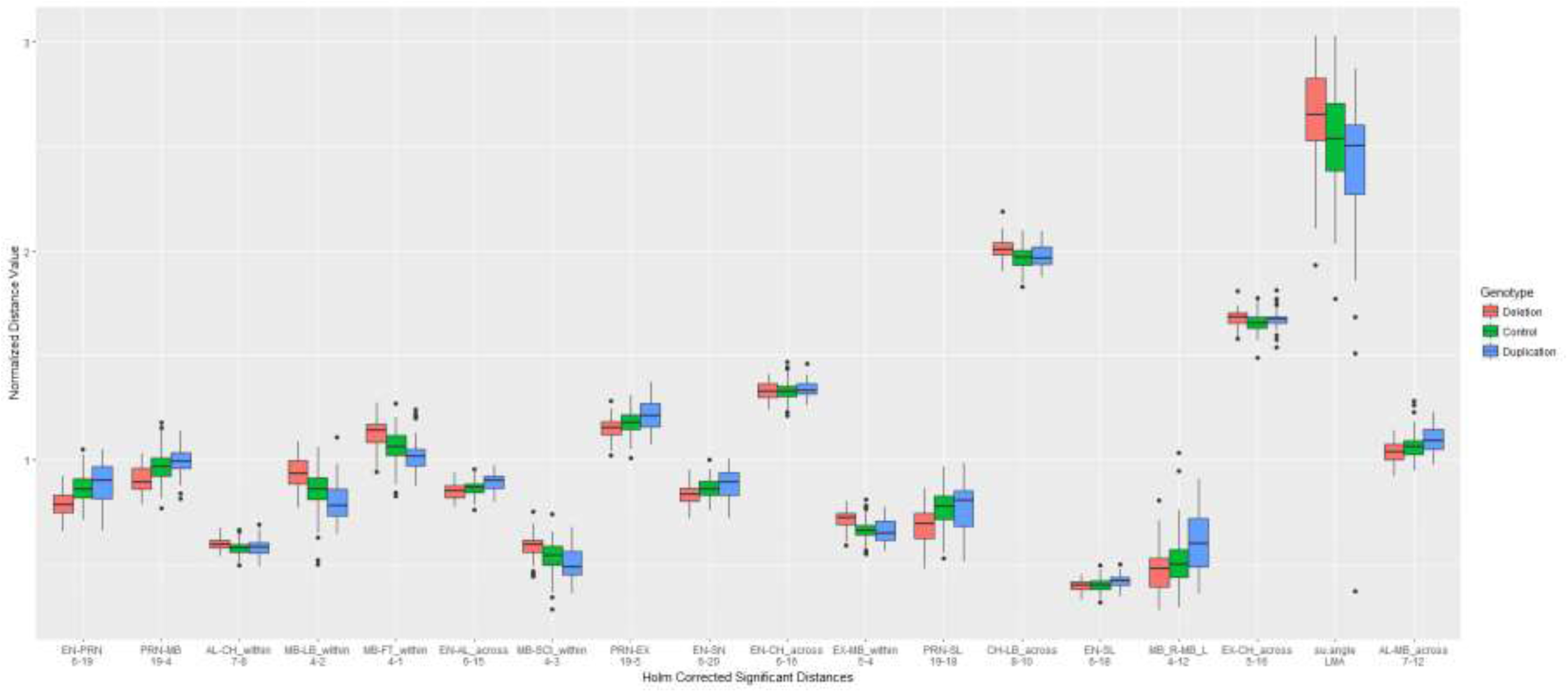
Craniofacial features that distinguish deletion, duplication and control groups. Eighteen craniofacial measures were differed significantly between CNV carriers and controls. Box plots illustrate the mean and standard error (whiskers) across the full dataset.

**Figure S3.**
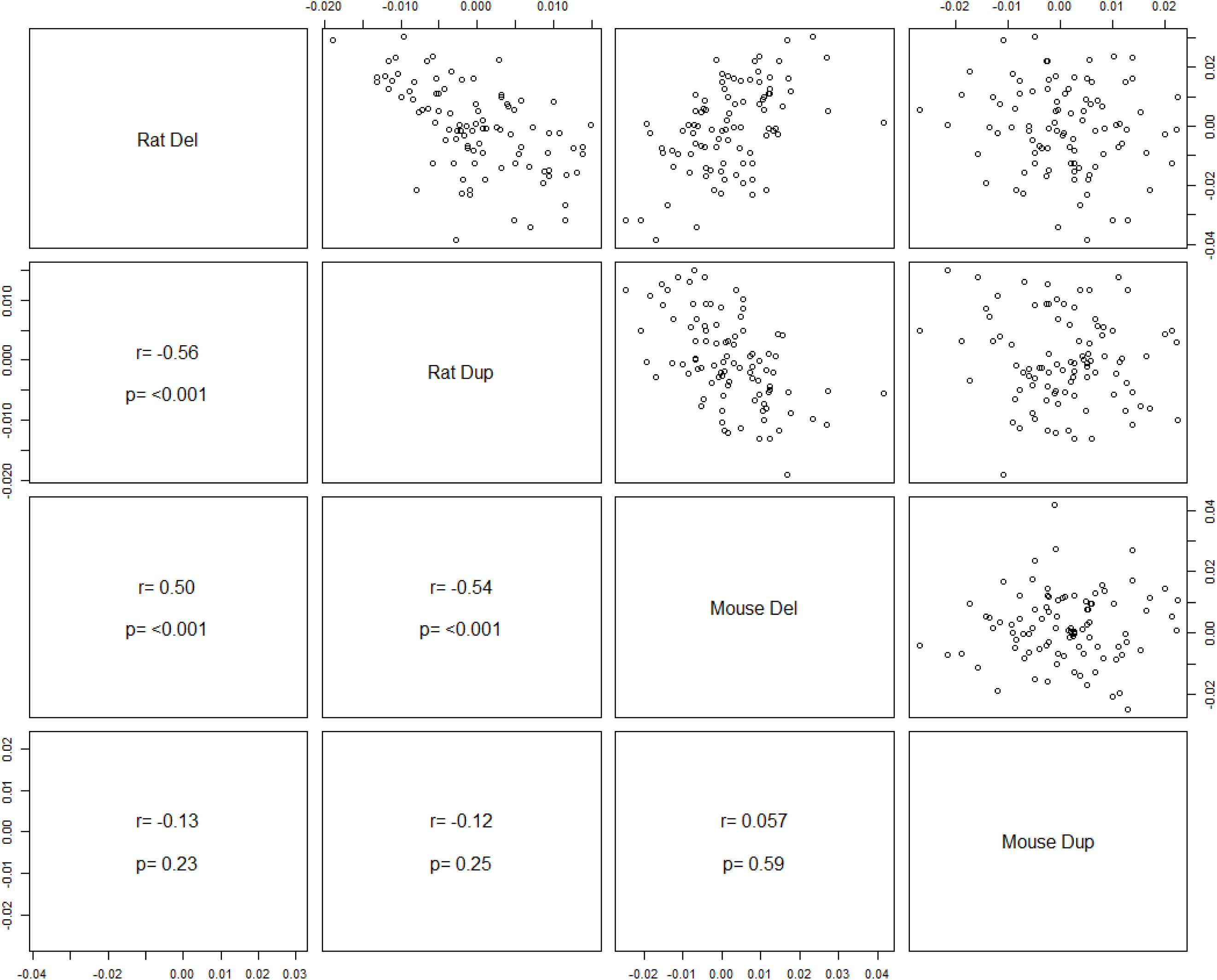
Comparison of the craniofacial effects of CNV in rat and mouse. Significant mirror effects of the deletion and duplication across all facial are observed in rat across all facial features. A significant correlation of effects was also observed between the rat and mouse models of the deletion. The effects of the duplication in mouse did not show a significant correlation with the other rodent models.

**Figure S4.**
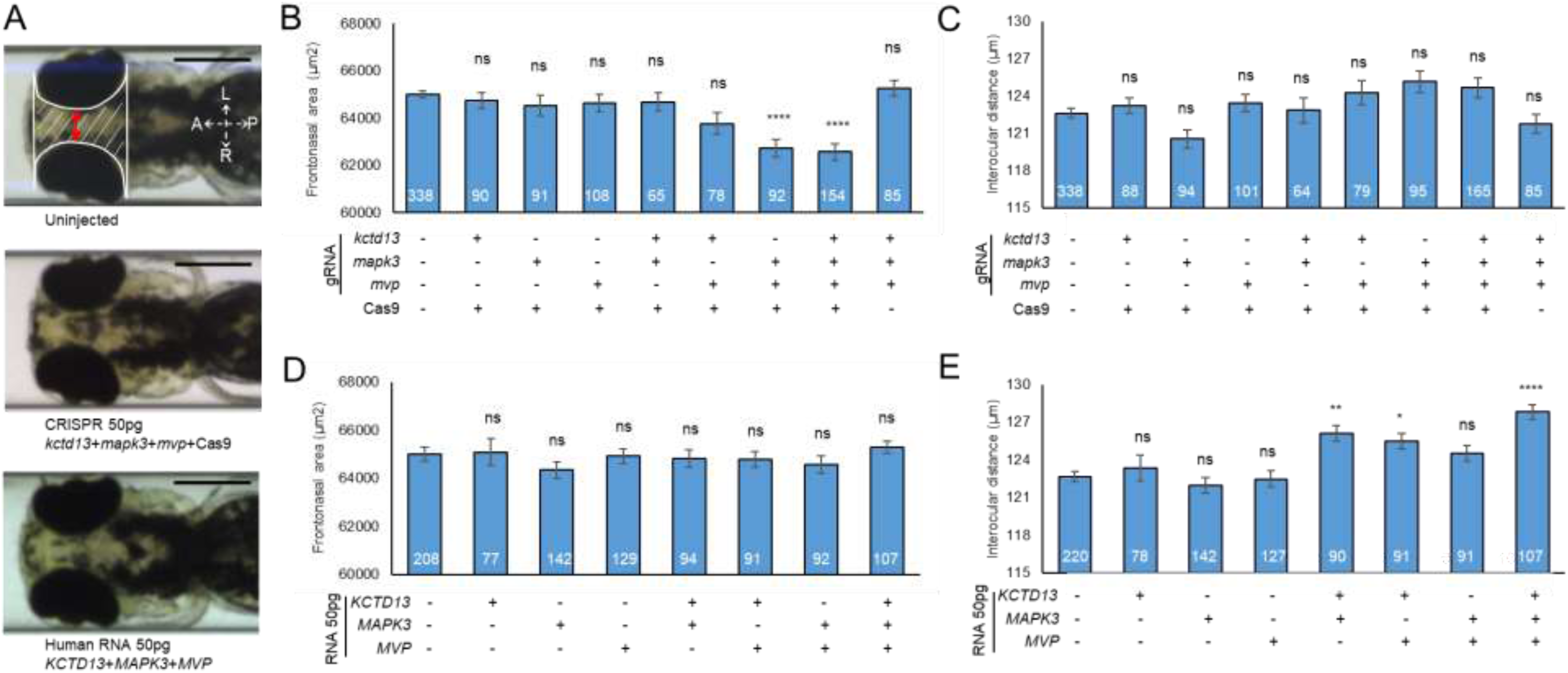
*KCTD13, MAPK3*, and *MVP* dose combinatorial suppression result in a decreased frontonasal area whereas similar overexpression results in an increased interocular distance. (**A**) Representative ventral images of *-1.4col1a1:egfp* zebrafish larvae at 3 days post-fertilization (dpf). Orientation arrows indicate anterior (A), posterior (P), left (L) and right (R). Area between the eyes: dashed white line. Interocular distance: red line. Scale bar, 200 µm. (**B, D**) Quantitative assessment of the frontonasal area. (**C, E**) Quantitative assessment of the interocular distance. The data are represented as the mean ±s.e.m.; ns=not significant; \**P*<0.05, \*\**P*<0.01 and \*\*\*\**P*<0.0001 vs uninjected controls. Tukey’s test was applied following a significant one-way ANOVA

**Figure S5.**
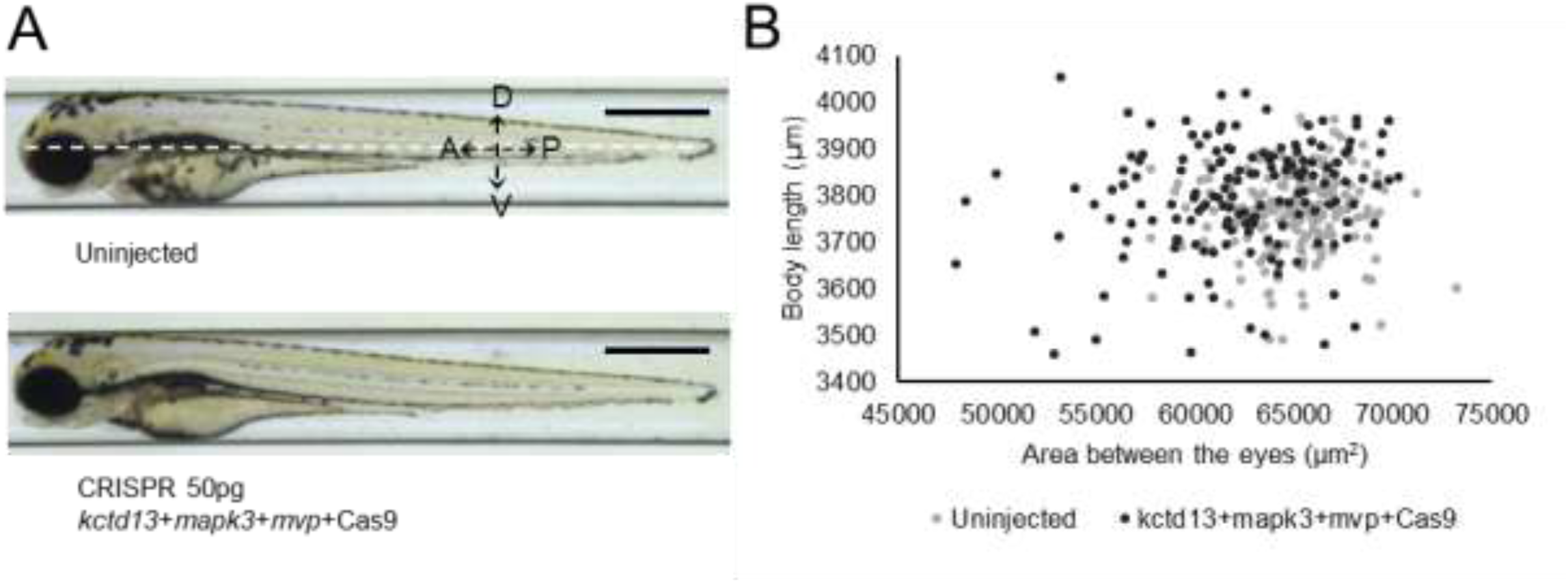
*kctd13, mapk3*, and *mvp* dose combinatorial suppression do not induce growth delay. (**A**) Representative lateral images of *-1.4col1a1:egfp* zebrafish larvae at 3 dpf. Measurement of the body length is shown with a white dashed line. Orientation arrows indicate anterior (A), posterior (P), dorsal (D) and ventral (V). Scale bar, 600 μm. (**B**) Scatterplot of all larvae; x-axis, area between the eyes; y-axis, body length. Each dot corresponds to one larva. F0 mutant larvae injected with equivalent amounts of *kctd13, mapk3*, and *mvp* guide do not show a reduction of the body length.

**Figure S6.**
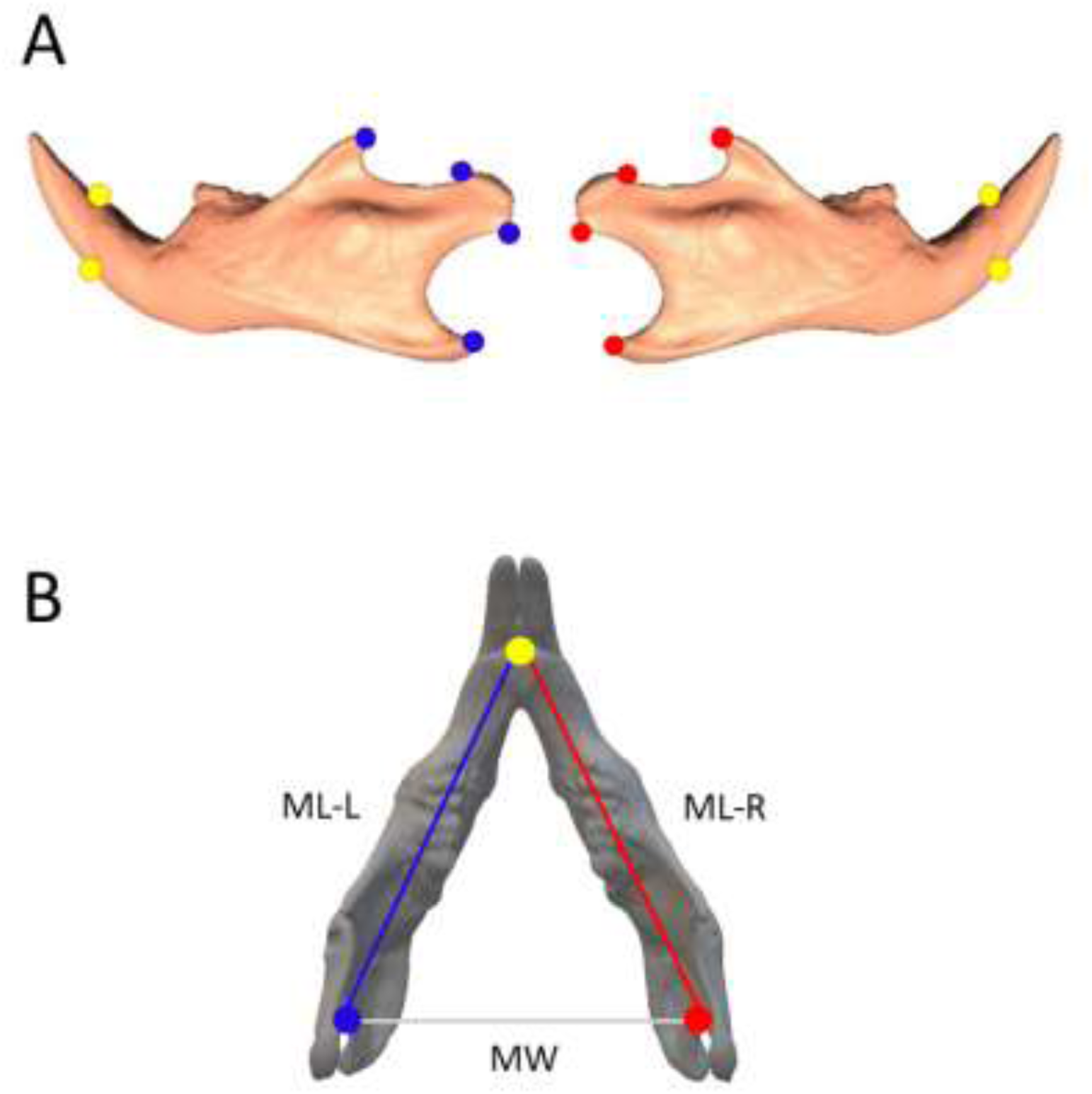
Landmarks that were used to determine the length and width of the mouse mandible. **(A)** On each jaw bone, two posterior landmarks were placed above and below the incisors (yellow) and four anterior landmarks were placed along the ramus (blue = left, red = right). **(B)** Landmarks were grouped by color and the centroid of each group was calculated from X,Y,Z coordinates. The distances between the three centroids was then used to determine the mandibular width (MW) and the length of the left (ML-L) and right (ML-R) mandible.

**Figure S7.**
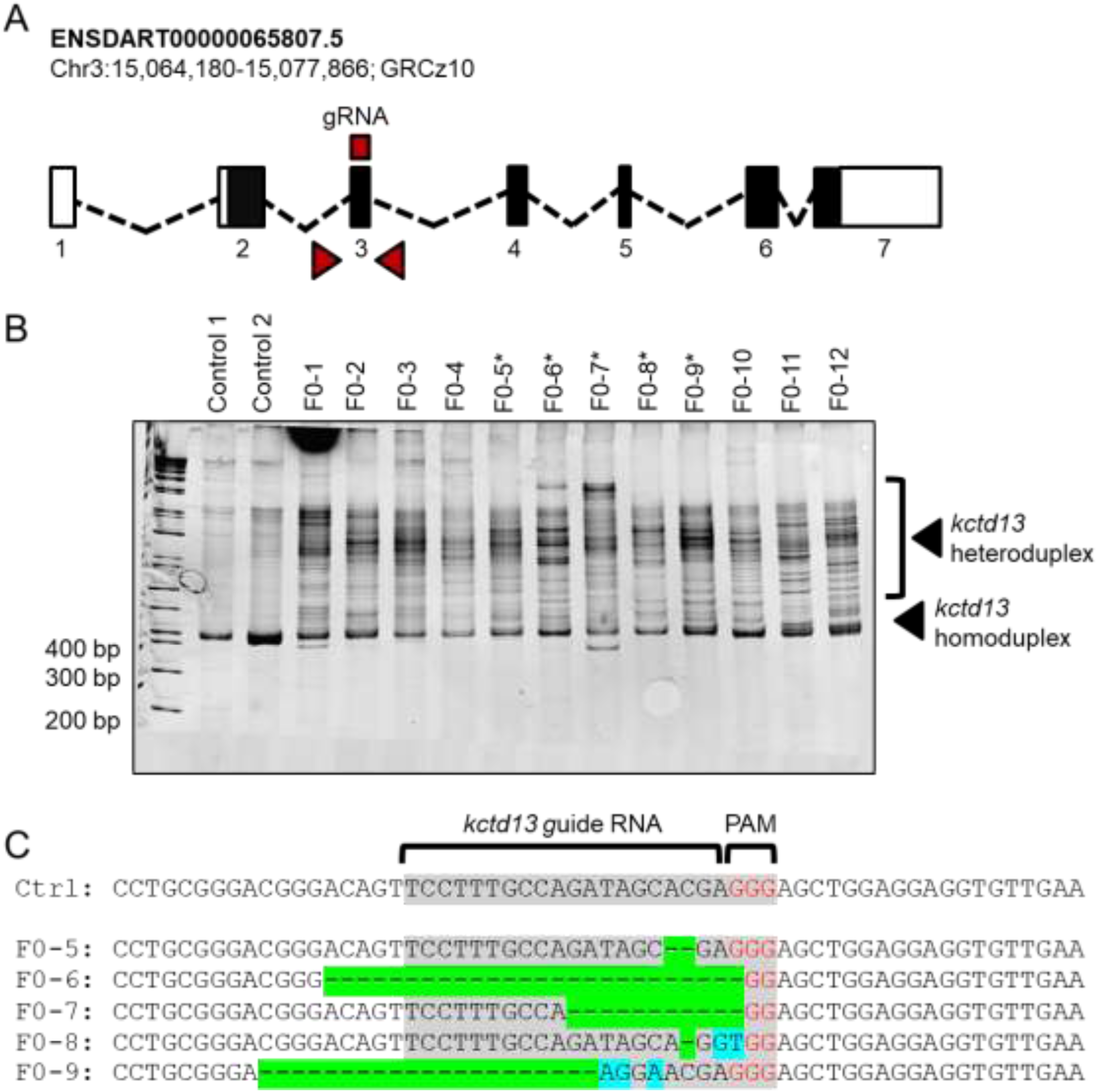
Genome editing of *kctd13* using CRIPSR/Cas9 to generate F0 zebrafish mutants. (**A**) Schematic of the *kctd13* ortholog in zebrafish. The locus is shown with exons (black boxes); untranslated regions (white boxes); introns (dashed lines); guide (g)RNA target site and primers used to generate PCR products shown in panel B (red box and triangles, respectively). (**B**) Assessment of genome-editing efficiency using polyacrylamide gel electrophoresis (PAGE). Genomic DNA was extracted from single embryos at 2 dpf, and PRC amplified. PCR products were denatured, reannealed slowly and migrated on a 20% polyacrylamide gel. All twelve F0 embryos displayed heteroduplexes not present in two uninjected controls. Asterisks (*) indicate embryos assessed for percent mosaicism with PCR8/GW/TOPO-TA cloning and Sanger sequencing of individual clones. (**C**) Representative sequence alignments showing the most common targeting events for each embryo. To estimate percent mosaicism, one control and five F0 embryos were assessed (n=9-12 clones/embryo); all FO clones harbored deletions (green) and some clones harbored insertions (blue), suggesting ∼100% efficiency. gRNA sequence (gray) and protospacer adjacent motif (PAM, red) are shown.

**Figure S8.**
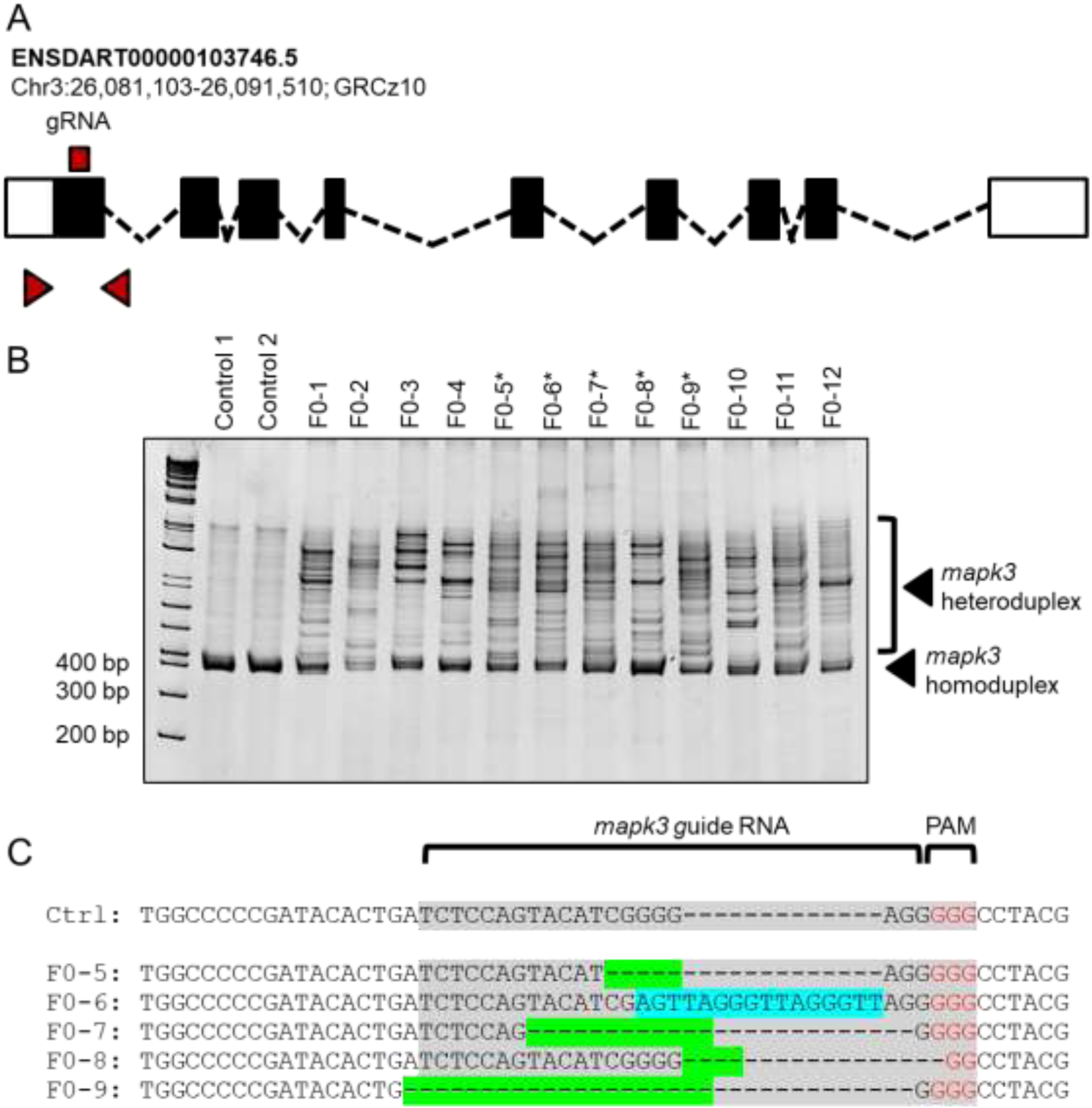
Genome editing of *mapk3* using CRIPSR/Cas9 to generate F0 zebrafish mutants. (**A, B**): see Figure S2. (**C**) A majority of F0 clones harbored deletions (green) and some clones harbored insertions (blue), suggesting ∼97% mosaicism.

**Figure S9.**
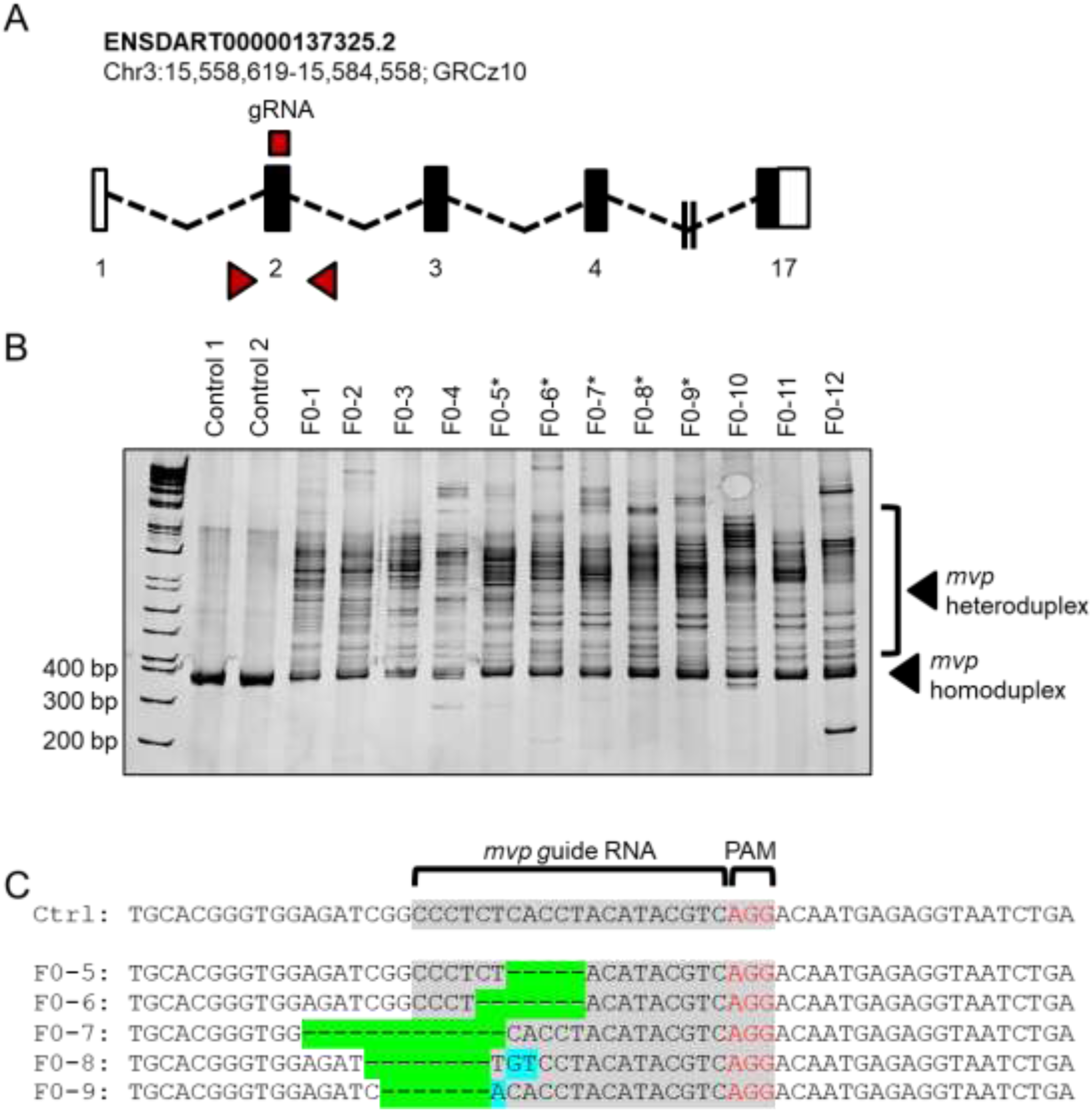
Genome editing of *mvp* using CRIPSR/Cas9 to generate F0 zebrafish mutants. (**A, B**): see Figure S2. (**C**) A majority of F0 clones harbored deletions (green) and some clones harbored insertions (blue), suggesting ∼96% mosaicism.

**Figure S10.**
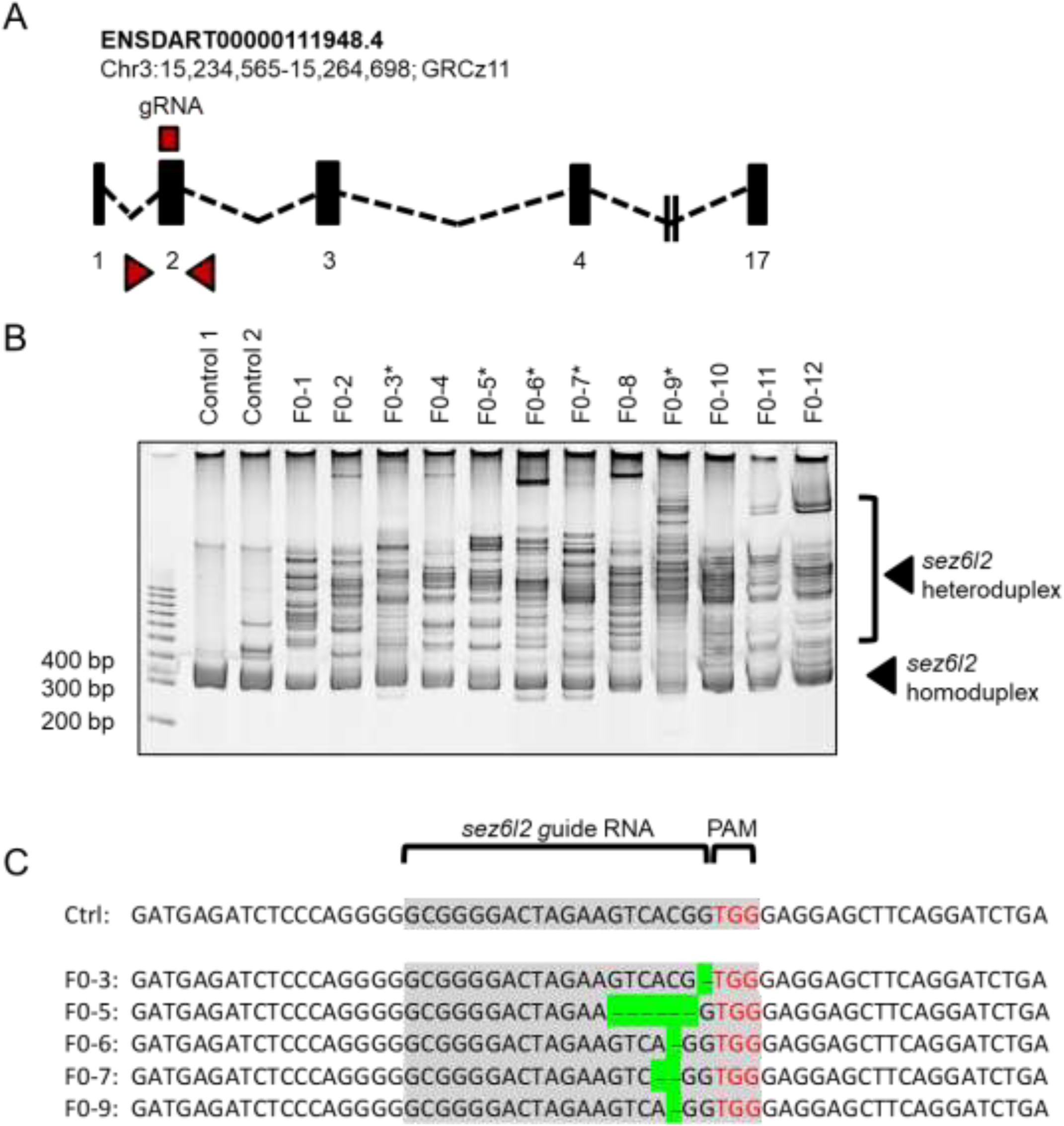
Genome editing of *sez6l2* using CRIPSR/Cas9 to generate F0 zebrafish mutants. (**A, B**): see Figure S2. (**C**) A majority of F0 clones harbored deletions (green), suggesting ∼96% mosaicism.

**Figure S11.**
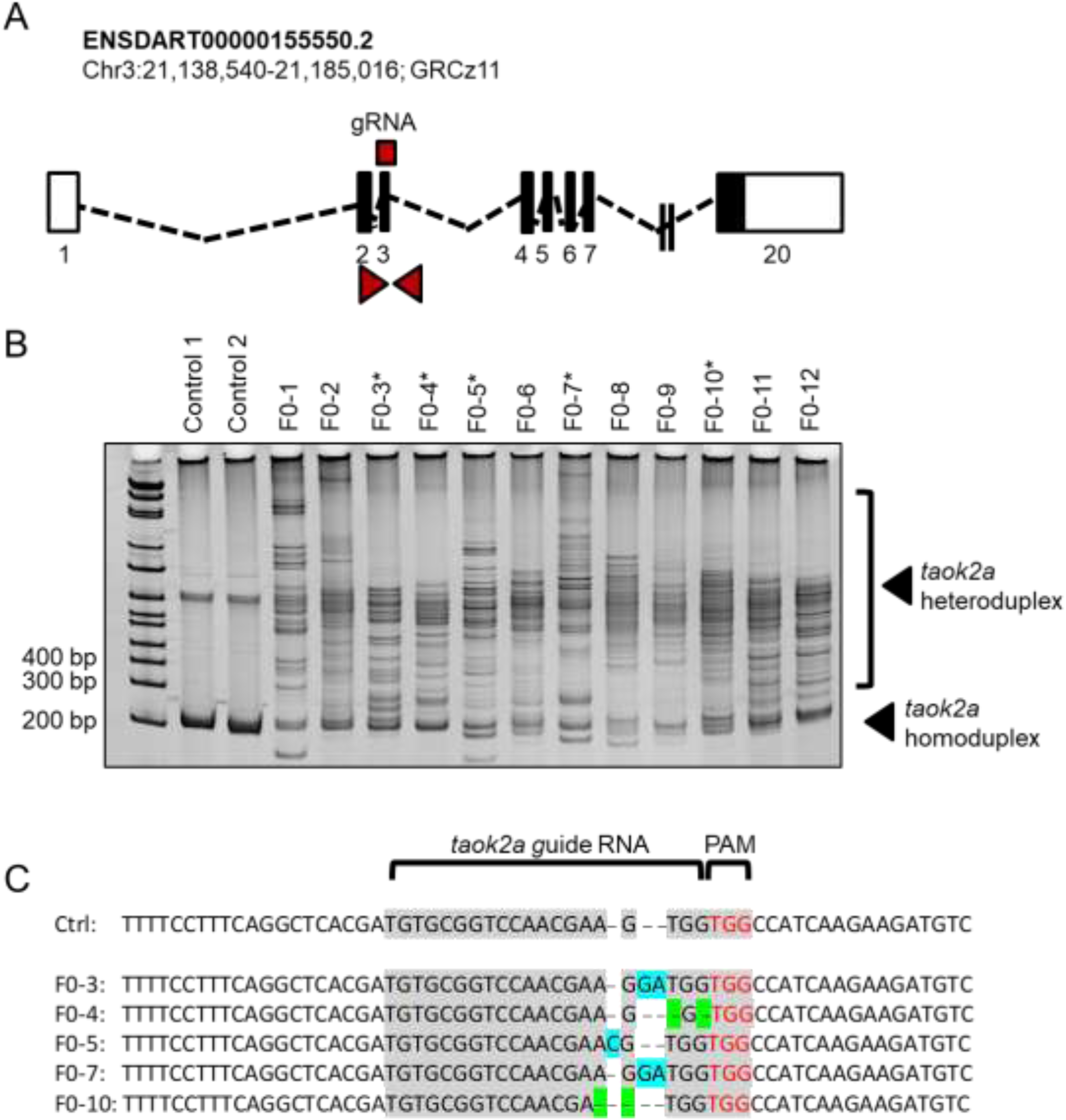
Genome editing of *taok2a* using CRIPSR/Cas9 to generate F0 zebrafish mutants. (**A, B**): see Figure S2. (**C**) Some F0 clones harbored deletions (green) and some clones harbored insertions (blue), suggesting ∼94% mosaicism.

**Figure S12.**
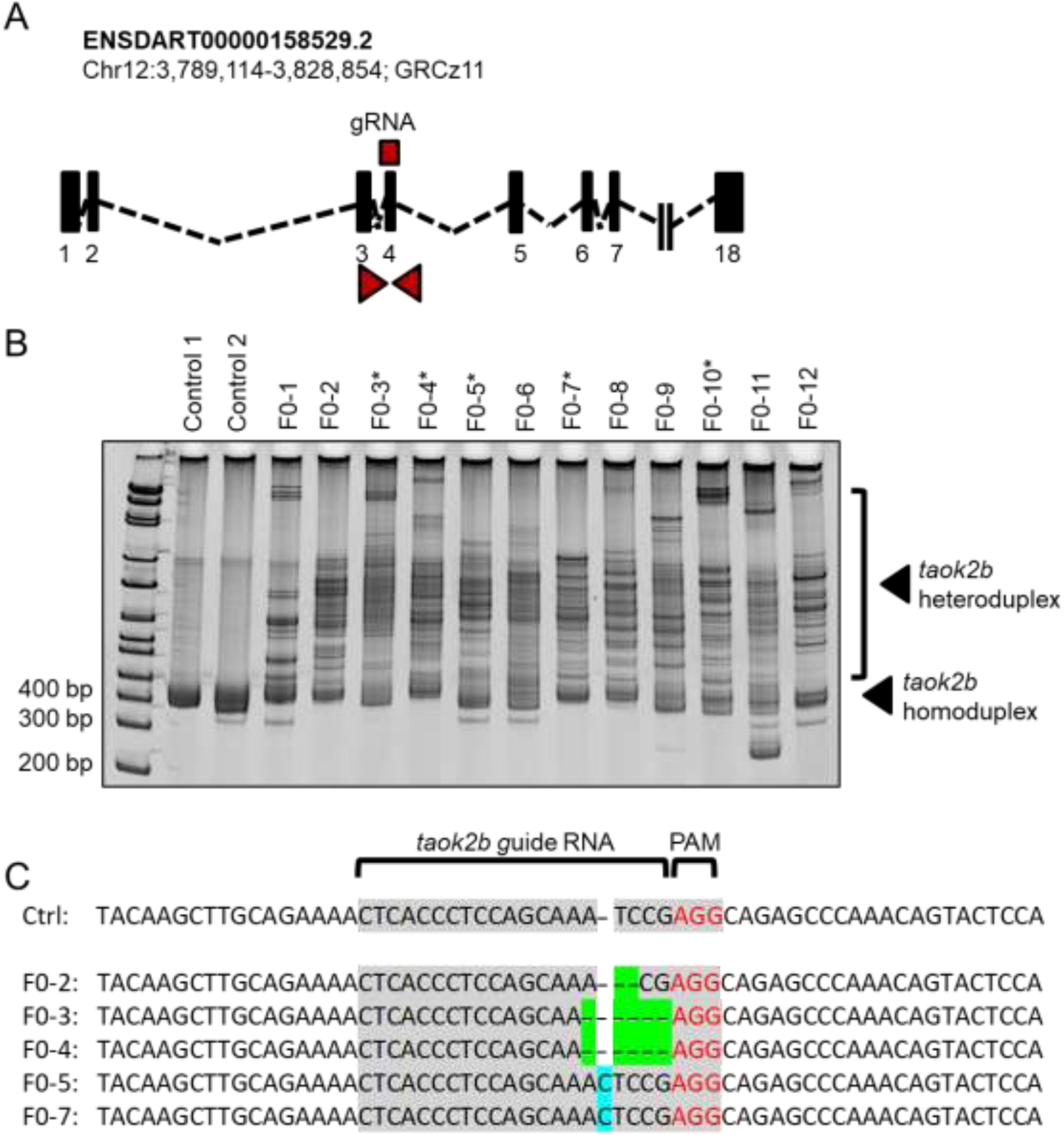
Genome editing of *taok2b* using CRIPSR/Cas9 to generate F0 zebrafish mutants. (**A, B**): see Figure X. (**C**) Some F0 clones harbored deletions (green) and some clones harbored insertions (blue), suggesting ∼89% mosaicism.

## Supplementary Methods

### Study Sample

Subjects were recruited in conjunction with the Simons VIP study (Consortium). 3D morphometric facial imaging was performed on a subset (N = 359) of subjects with 16p11.2 duplications or deletions from the Simons VIP cohort at 3 sites (UW, Harvard, Baylor) using the 3DMD 3-pod camera system. Data analysis was restricted to subjects of European ancestry older than 3 years of age. Additional subjects were excluded due to facial hair, image quality or landmark visibility (i.e. obscured by clothing, hair or makeup). The final dataset (N = 228) included 45 deletions, 44 duplications and 139 familial non-carrier controls (**Table S1**).

### 3DMD morphometric imaging

Images were acquired using the 3dMDtrio system (http://www.3dmd.com/3dmd-systems/#trio). Images were landmarked according to Farkas standards (Farkas) blind to genotype. In addition to standard Farkas landmarks, additional landmarks were placed to capture frontal dimensions including landmarks 2 (lateral brow) and 4 (medial brow). A total of 24 landmarks were placed (**Fig. 1A**). All Landmarking was performed by a single analyst (S.T.) blind to genotype. To confirm a high reliability of our manual landmarking, an independent set of landmarks was generated by a second analyst (T.P) and reproducibility was determined by the intraclass correlation of the two sets of distances measures (127 distances per set), assuming random effects for both subjects and analysts. The median ICC across all distances was 0.89; 87% (110/127) of distances had ICC > 0.7; and 98% (125 out of 127) show no significant difference between two analysts (p<0.05).

#### Analysis

Quantitative measurement of all pairwise distances between 24 landmarks were calculated using the 3dMDvultus – Analysis software, version 2.5.0.1 (3dMD.com). Symmetric distances were averaged, yielding 156 facial distance measurements. Each distance was normalized to the overall size of the individual’s face, by dividing by the geometric mean of the 156 distances for that individual. Angular measurements of the nose and chin, the nasomental (NMA) and labiomental (LMA) angles respectively, were calculated by triangulating the relevant landmarks A series of linear mixed-effects models, using package lme4 in R (version 3.4.1), was used to separately test for the effect of deletion and of duplication on each angle or normalized facial distance. Each model controlled for fixed effects of age, head circumference, body mass index (BMI), sex, and ancestry principal components, with a random intercept allowed to account for within-family correlation. Interaction between genotype and sex was included if significant at 5% level. Significant differences according to genotype were determined by a likelihood-ratio test at a family-wise error rate of 5% using Holm’s correction (Holm).

### Controlling for variation in ancestry

All subjects were of European ancestry, however regional genetic differences could still explain variation in facial traits. We controlled for ancestry using principal components derived from genetic data subjects. Ancestry principal components were obtained on 213 subjects from Illumina SNP genotype data (Illumina HumanOmniExpress v.1 and v.2) available from the Simons VIP study (https://www.sfari.org/resource/simons-vip/). Missing data on 38 subjects was imputed by using PCs from a sibling nearest in age or a randomly selected parent. After imputation, 15 subjects from two different families were still missing data, 9 of which were familial controls and 6 were duplications. Analyses were performed with and without including the ancestry principal components as a sensitivity analysis, and the results were very similar. Significant correlation was observed between one of the first two principal components and 7 of the 160 craniofacial distances.

### Generating 3D models by averaging faces of deletion, duplication and control subjects

To visualize the respective facial gestalts of controls, deletion carriers and duplication carriers, a 3D-model of each was generated by averaging of the surface topography of faces from multiple subjects using the 3dMDvultus – Analysis software, version 2.5.0.1 (3dMD.com). To maximize the number of unrelated subjects that were closely matched in age within each group, selection criteria for averaging of faces differed slightly from that of the overall dataset. Only unrelated individuals were included, additional subjects were removed due to image quality (gaps in the surface topography), and the requirement for landmark visibility was relaxed, allowing for frontotemporal landmarks (landmarks 1 and 9, see **Fig. 1**) to be covered in some cases by hair or headwear. We first restricted our 3D models to young adults (age 14) and older to avoid variability in facial features across development at young ages, and the sample was restricted to males (the largest group). Subjects consisted of 5 Deletion carriers (average age 25.5 years), 5 duplication carriers (36.5 years) and 10 controls (36.6 yrs). Four Landmarks (the Exocanthion, Glabella and Subnasale, Fig 1. Landmarks 5, 13, 17, 20) were placed manually, and the software’s average-face function was used to generate the average face. The surface property of the 3D image was then converted from a photographic image into a textured-model (**Fig. 1D**). Subsequently, to determine if similar facial gestalts are apparent for other demographics, additional 3D models were generated for children and females (**Fig. S1**). Subjects that were included in average face models are listed in **Table S3**.

### Linear Discriminate Analysis

Linear discriminate analysis (LDA) was performed on linear model residuals after adjusting for age, head circumference, body mass index (BMI), sex, and ancestry principal components, for both total subjects and subjects aged < 20 years. The 45 distances with FDR q value less than 0.05 were used. We used the function “lda” from the “MASS” package in R (version 3.4.1). Specificity and sensitivity were calculated based on LDA prediction with default values.

### Least absolute shrinkage and selection operator (LASSO) logistic regression

Generalized linear model with lasso was performed for both human subjects and mice. Distances with FDR q value less than 0.05 were used for human subjects, while distances with a statistically significant likelihood ratio test at p <.05 were used for mouse skulls. We used 10-fold cross validation for lasso with minimum deviance for human subjects, and lasso with minimum Akaike Information Criteria (AIC) for mouse skulls (Akaike). Specificity and sensitivity were calculated based on lasso selected models. Packages “glmnet” and “glmpath” from R (version 3.4.1) were used.

### Description of rat and mouse cohorts

Craniofacial structure was analyzed for rodent models of 16p11.2 Deletion and Duplication in rat and mouse. Rat deletion and duplication models were generated by CRISPR/Cas9 genome editing of Sprague Dawley line (Charles River Laboratory, Oncins, France). Briefly a deletion of 483,122 bp located at positions chr1:198,100,544-198,583,667 (RatRnor_6.0) and a duplication of the interval from chr1:198,100,545-198,583,458 (RatRnor_6.0), corresponding to the 16p11.2 homologous region of the rat genome, were obtained. For the genotyping, primer pairs were designed for the Del, Dup and an internal control alleles (Primers Del: rHamont99For: GGGCTGGCAGACTTGAA rHavalB284Rev: GTGCCACGATCAGCAG; Primers Dup: rHamont99Rev: CGCTTTGATGCCCACTA; rHavalB84For: AGCTGTGATCCTCTGGTT; Primers for internal control: rAnks3-205For: CCCCAGCCTCCCACTTGTC, rAnks3-205Rev: AGGATGACTGAAATTGGTGGAC) to amplify specific PCR fragments (Del: 290bp, Dup: 500bp, internal control 205bp) using standard conditions (Roche, 60°C for primer hybridation). A cohort of 75 rats was bred for craniofacial analysis, which included 23 *Del/+* (9 male and 14 female), 26 *Dup/+* (13 male and 13 female) and 26 *+/+* siblings (13 male and 13 female). The mouse models of 16p11.2 used in this study consisted of deletion (*Del/+*) or duplication (*Dup/+*) of the *Sult1a1-Spn* genetic interval (Arbogast et al.) Lines were maintained on a pure C57BL/6N C3B genetic background. A mouse cohort was bred including 36 females at 13 weeks of age, including 10 *Del/+* and 10 *+/+* littermates, and 8 *Dup/+* and 8 *+/+* littermates.

### Rodent skull Imaging, Landmarking, Data Processing

For both rat and mouse cohorts, images of the dorsal skulls were captured using a microCT imaging system (Quantum GX, Perkin Elmer, France). For rats, an image was acquired for the complete skull. For mice, images of the dorsal skull and lower jaw of each animal were acquired separately as part of a previous study (Arbogast et al.). Nineteen landmarks were placed representing the frontal, nasal and maxillary regions, and all pairwise distances between landmarks were normalized to the geometric mean. In addition the mandibular length (ML) and width (MW) were determined by first determining the centroid of multiple landmarks at the lower incisors and the left and right ramus (**Fig. S6**), and then determining distances between the three centroids. Symmetric distances of the skull and mandible were averaged.

Differences in facial features between deletion and control lines and differences between duplication and control lines were tested with univariate linear models. We tested all 91 distances on the dorsal skull and two distances on the mandible. Effects that were significant at an FDR of 5% were identified. In addition, as we did previously in human, we identified a set of features that distinguish CNV models from controls by performing least absolute shrinkage and selection operator (LASSO) based on all univariate significant distances by generalized linear model, with AIC as the criteria rule.

### mRNA overexpression and CRISPR/Cas9 genome editing in zebrafish embryos

Zebrafish embryos were obtained from natural matings of heterozygous *-1.4col1a1:egfp* transgenic adults maintained on an AB background (Kague et al., 2012). To model the 16p11.2 BP4-BP5 duplication, we overexpressed individually each gene of the region (see Fig. 4D). We linearized pCS2+ constructs (Golzio et al., 2012) and transcribed human mRNA using the mMessage mMachine SP6 Transcription Kit (Ambion). All RNAs were injected into the yolk of the embryo at the 1-to 4-cell stage at 50, 25, or 12.5 pg doses (1 nl/injection). To investigate specific gene interactions that have been reported previously (Golzio et al.), *KCTD13, MAPK3,* and *MVP* mRNAs were tested in combinations of two or three. Two way and three way gene interaction models were fitted to test the synergy effect from double-hit or triple-hit groups. Packages “multcomp” from R (version 3.4.1) was used.

CRISPR/Cas9 genome editing was performed as a model of the reciprocal deletion. We used CHOPCHOP(Labun et al., 2016) to identify guide (g)RNAs targeting coding sequence within *kctd13, mapk3, mvp, sez6l2, taok2a*, and *taok2b* (**Table S7**). gRNAs were transcribed *in vitro* using the GeneArt precision gRNA synthesis kit (ThermoFisher) according to the manufacturer’s instructions; 1 nl of injection cocktail containing 50 pg/nl gRNA and 200 pg/nl Cas9 protein (PNA Bio) was injected into the cell of embryos at the 1-cell stage. To determine targeting efficiency in founder (F0) mutants, we extracted genomic DNA from 2 day post-fertilization (dpf) embryos and PCR amplified the region flanking the gRNA target site. PCR products were denatured, reannealed slowly and separated on a 20% TBE 1.0-mm precast polyacrylamide gel (ThermoFisher), which was then incubated in ethidium bromide and imaged on a ChemiDoc system (Bio-Rad) to visualize hetero- and homoduplexes. To estimate the percentage of mosaicism of F0 mutants (*n* = 5/condition), PCR products were gel purified (Qiagen), and cloned into a pCR8/GW/TOPO-TA vector (Thermo Fisher). Plasmid was prepped from individual colonies (*n* = 9–12 colonies/embryo) and Sanger sequenced according to standard procedures.

### Automated zebrafish imaging

Larvae were maintained under standard conditions at 28.5°C until 3 dpf and were positioned and imaged live as described (Isrie et al., 2015). Automated imaging was conducted with an AxioScope.A1 microscope and Axiocam 503 monochromatic camera facilitated by Zen Pro software (Zeiss), to capture dorsal images of GFP signal. Larval batches were positioned and imaged live using the Vertebrate Automated Screening Technology (VAST; software version 1.2.5.4; Union Biometrica) BioImager. Larvae from each experimental condition were anesthetized with 0.2 mg/mL Tricaine prior to being loaded into the sample reservoir. Dorsal and lateral image templates of uninjected controls and experimental larvae were created and we acquired images at a >70% minimum similarity for the pattern-recognition algorithms. Larvae were rotated to 180° to acquire ventral images via a 10x objective and fluorescent excitation at 470nm to detect GFP to capture fluorescent images of the pharyngeal skeleton. ImageJ software (NIH) was used to measure the angle of the ceratohyal cartilage. All experimental conditions were normalized to uninjected controls and set to 100 degrees. Statistical comparisons were performed using one-way ANOVA with Tukey’s test (GraphPad Prism).

### Examining levels of gene expression during murine craniofacial development

We examined whether the significant effects of seven genes (*SPN, C16orf54, SEZ6L2, ASPHD1, TAOK2, INO80E and FAM57B*) on shape of the mandible could be attributable to the differential regulation of these genes. A published dataset was obtained consisting of Affymetrix gene expression analysis of the major craniofacial processes of the developing mouse embryo (E10.5-E12.5) (Hooper et al.) (accession # FB-STU-201-0001, Facebase.org). Samples included mesenchymal and ectodermal cells of the frontonasal and mandibular processes of embryos at E10.5, E11.5 and E12.5 in triplicate, and samples of the maxillary process at E11.5 and E12.5. The basal expression levels of the seven genes was compared to the levels of other in each structure was determined by averaging the expression values across replicates and embryonic stages. Results show the expression levels of all three genes to be consistent across cell types and structures of the face (**Fig S9**).

